# Deep mutationally scanned (DMS) CHIKV E3/E2 virus library maps viral amino acid preferences and predicts viral escape mutants of neutralizing CHIKV antibodies

**DOI:** 10.1101/2024.12.04.626854

**Authors:** Megan M. Stumpf, Tonya Brunetti, Bennett J. Davenport, Mary K. McCarthy, Thomas E. Morrison

## Abstract

As outbreaks of chikungunya virus (CHIKV), a mosquito-borne alphavirus, continue to present public health challenges, additional research is needed to generate protective and safe vaccines and effective therapeutics. Prior research has established a role for antibodies in mediating protection against CHIKV infection, and the early appearance of CHIKV-specific IgG or IgG neutralizing antibodies protects against progression to chronic CHIKV disease in humans. However, the importance of epitope specificity for these protective antibodies and how skewed responses contribute to development of acute and chronic CHIKV-associated joint disease remains poorly understood. Here, we describe the deep mutational scanning of one of the dominant targets of neutralizing antibodies during CHIKV infection, the E3/E2 (also known as p62) glycoprotein complex, to simultaneously test thousands of p62 mutants against selective pressures of interest in a high throughput manner. Characterization of the virus library revealed achievement of high diversity while also selecting out non-functional virus variants. Furthermore, this study provides evidence that this virus library system can comprehensively map sites critical for the neutralization function of antibodies of both known and unknown p62 domain specificities.

**IMPORTANCE:** Chikungunya virus (CHIKV) is a mosquito-borne alphavirus and re-emerging pathogen of global health concern that causes fever and debilitating acute and chronic joint pain. Prior studies established a critical role for antibodies in protection against CHIKV infection, however, the importance of different antibody epitopes engaged on the virus for these functions is poorly understood. Here, we describe the generation of a high-throughput, functional virus library capable of simultaneously identifying critical neutralization sites for multiple antibodies. We find that, through generation of the virus library, the plasticity of the viral glycoproteins tested varied by subdomains. Additionally, our study revealed new sites in the major CHIKV surface glycoprotein important for neutralization by previously proposed therapeutic monoclonal antibodies. Overall, this study describes a new tool that can be used to better understand antibody responses associated with distinct CHIKV infection outcomes and could contribute to the development of efficacious vaccines and antibody-based therapeutics.

## INTRODUCTION

Chikungunya virus (CHIKV), a mosquito-borne alphavirus, causes fever, severe acute and chronic joint pain and inflammation, and is a re-emerging public health threat (1–5). Moreover, CHIKV infection can be fatal; the case-fatality ratio (CFR) has been estimated between 1-2.2 deaths per 1,000 cases in both Asia and the Americas (6, 7). Outbreaks continue to increase in frequency and size, due in part to adaptive mutations in the viral surface E1 and E2 glycoproteins allowing for improved transmission efficiency by *Aedes albopictus* and *Aedes aegypti* mosquitoes (8, 9). Prior research established a role for anti-CHIKV antibodies (Abs) in control of CHIKV infection, and human studies suggest that the early appearance of CHIKV-specific IgG or IgG neutralizing Abs (nAbs) protects against progression to chronic CHIKV disease (10–16). However, the importance of epitope specificity of protective Abs and how skewed responses contribute to resolution of symptoms or progression to chronic disease remains poorly understood.

CHIKV is an enveloped, positive-sense RNA virus with a ∼12 kb genomic RNA that encodes two open reading frames (ORFs). The first ORF encodes a non-structural (ns) polyprotein that is processed to produce four ns proteins (nsP1-4) that mediate RNA synthesis. The second ORF encodes the viral structural proteins capsid-p62-6K/TF-E1. E1 and p62 co-translationally associate within the ER and are glycosylated (17). Within the secretory pathway, p62 is cleaved by furin into the mature E2 and E3 glycoproteins (18). Although the E3 glycoprotein is cleaved during the maturation process, for some alphaviruses, including CHIKV, it can remain bound to the mature E2–E1 heterodimer (19, 20). E2 and E1 heterodimers form 80 trimeric spikes on the viral surface. The E2 protein is comprised of three domains (A, B and C), with domain A positioned towards the center of the trimeric spike, domain B on the tip of the spike, and domain C next to the viral membrane. E1 is a class II fusion protein with three domains termed I, II and III. E2 and E1 are the dominant targets for nAbs (21–27).

Here, we deep mutationally scanned (DMS) (28) the CHIKV p62 glycoprotein to simultaneously test thousands of p62 mutants against selective pressures of interest in a high throughput manner. DMS of viral proteins has been employed for several viruses, including HIV, SARS-CoV-2, hepatitis C virus (HCV), Mayaro virus, Zika virus, and others (28–36). This library generation method has a wide variety of applications, ranging from the evaluation of therapeutic mAbs and their epitope sites to informing phylogenetic viral evolution models (28, 37–41). Certain DMS methods involve the use of replication incompetent systems to avoid the need for higher biosafety level facilities (34, 35, 42, 43). Because CHIKV p62 (E2/E3) is involved in cell entry and egress, we generated a full-length CHIKV plasmid system to elucidate how viral fitness may be affected by certain mutations.

In this study, we evaluated the diversity of the CHIKV-p62-DMS library and inferred the mutational tolerance of the CHIKV p62 structural glycoprotein complex. We then utilized two well-characterized CHIKV mouse mAbs (CHK-152 and CHK-265) (21–26) to verify our library’s capacity to identify important functional sites for nAbs and ability to add to our current knowledge of mutations that can alter mAb neutralizing activity. In addition, we tested the library’s capacity to map sites that influence the neutralizing activity of a mAb whose precise target is undefined (CHK-11) (21). Results from our characterization of the CHIKV-p62-DMS virus library revealed high diversity while also selecting out non-functional virus variants. Furthermore, our data provides evidence that this virus library system can comprehensively identify sites that alter neutralization by mAbs of both known and unknown p62 domain specificities.

## RESULTS

### Characterization of the CHIKV-p62-DMS Virus Library Reveals High Diversity in mutDNA and Functional Variant Selection in mutVirus.p2

NAbs against CHIKV target the E2 and E1 surface glycoproteins (21–27), but little is known about the role of E3 in the nAb response. For these reasons, we deep mutationally scanned the p62 ectodomain to generate a library virus that could be used to identify mutations in E3/E2 that impact Ab-mediated inhibition of CHIKV infection. The CHIKV-p62-DMS virus library was generated in the context of a pCHIKV-CMV-mKate plasmid (**Figure S1A;** see **Materials and Methods**) (44). The plasmid library (mutDNA) was transfected into HEK293 cells, and virus-containing cell culture supernatants were collected at 48 hours post-transfection (hpt) to generate the first virus library (mutVirus.p1). To select the virus library for functional variants, Vero cells were inoculated with mutVirus.p1 (MOI of 0.01 PFU/cell) and virus-containing cell culture supernatants were collected at 48 hours post-infection (hpi) to generate mutVirus.p2 (**Figure 1A**). Vero cells were chosen for library virus generation because these cells are commonly used for monoclonal Ab (mAb) epitope mapping studies and neutralization assays (21–23, 26), allowing us to compare our findings with previously published contacts or escape residues. Two replicate libraries were independently generated, and both libraries were mutagenized to similar extents (see **Data and Code Availability** section).

**Figure 1.**
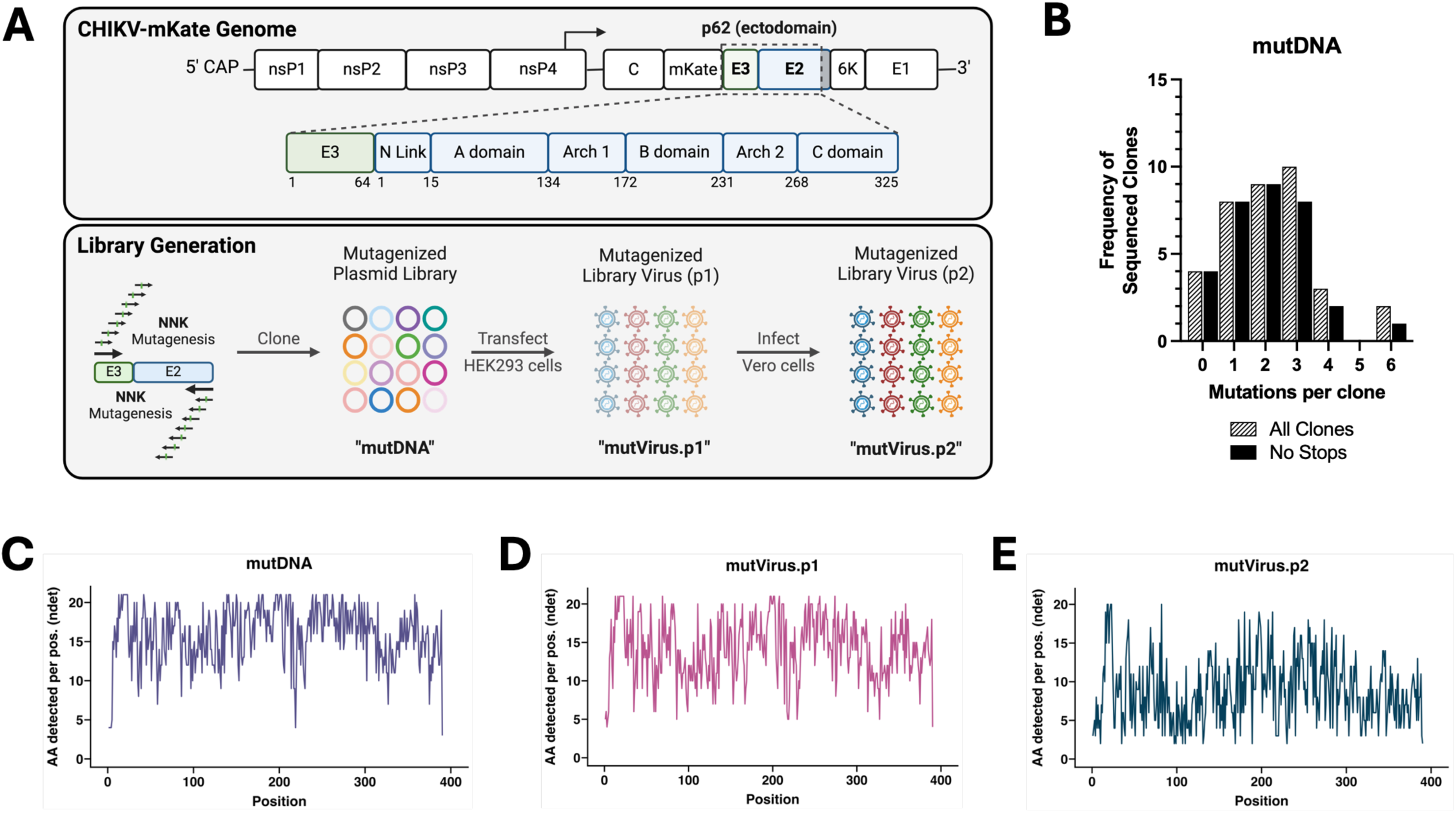
Generation of a deep mutationally scanned CHIKV p62 full-length virus library. (**A**) Schematic of the CHIKV genome (box indicates mutagenized region), procedure for generation of the mutagenized libraries, and naming scheme for each of the library iterations. (**B**) Sangar sequencing results for individual plasmid DNA clones. The lined black bars represent the number of nonsynonymous mutations per full-length CHIKV p62 clone (Sangar primers available in **Materials and Methods**). The solid black bars exclude any sequences containing stop codons. Average number of mutations per clone for the lined and solid black bars are 2.2 and 2.0, respectively. Distribution of these mutations across the CHIKV p62 region are detailed in **Figure S1C**. (**C-E**) Following deep sequencing, total number of detected amino acids per codon position (‘ndet’), for the (C) mutDNA, (D) mutVirus.p1, and (E) mutVirus.p2 libraries are plotted for the entire mutagenized CHIKV p62 region. Results for the wtDNA sequencing control are shown in **Figure S1D**.

To evaluate the efficiency of the mutagenesis, plasmid DNA isolated from 40 bacterial colonies, out of an estimated 2.1 x 10^4^ total, were Sangar sequenced to identify nonsynonymous mutations in the mutagenized p62 fragment (**Figure 1B**). 4/40 (10%) clones either lacked an insert from the ligation reaction or were bacterial contaminants. The average number of nonsynonymous and synonymous mutations in the 36 remaining clones was 2.2 and 0.08, respectively (**Figure 1B and Figure S1B**). 4/36 clones (11%) were eliminated from the analysis due to the presence of a stop codon, decreasing the average number of nonsynonymous mutations per functional clone to 2.0 (**Figure 1B**). The 36 clones with CHIKV p62 inserts were aligned to visualize the distribution of nonsynonymous mutations throughout the fragments, showing a relatively even spread among the sampled sequences (**Figure S1C**).

To assess library sequence diversity, RNA was isolated from mutVirus.p1 and mutVirus.p2, cDNA was generated, and the mutagenized p62 fragment was PCR-amplified. This same region also was PCR amplified from the mutDNA library and the WT pCHIKV-CMV-mKate plasmid, to account for PCR-based and sequencing-based errors, and the resulting amplicons were sequenced. Raw reads were cleaned and aligned to the pCHIKV-CMV-mKate reference genome, and codon frequencies and quality scores were calculated. Low-frequency and low-quality codons were filtered out of the data set and the number of remaining amino acids (AA) per codon position (‘*ndet*’) was analyzed. The average number of AA detected per position for the mutDNA, mutVirus.p1, and mutVirus.p2 libraries was 15.9, 14.2, and 8.9 AAs, respectively (**Figure 1C-E**). In contrast, the wtDNA sample had an average of 2.6 AAs detected per position (**Figure S1D**). These findings suggest that both a high degree of mutagenesis was achieved, and functional selection of the virus library occurred following passaging.

### Elevated Diversity in the E2 B Domain

To simplify the visualization of the deep sequencing data to the absence or presence of AAs above filters, we developed a new metric, ‘*ndet*’ (see **Materials and Methods**). This avoids confusion with large-effect sizes on rare mutants in logoplots that did not correspond to the frequency of that mutant in the library. To do this, we developed an alternative software package called *megaLogo* (https://github.com/meganstumpf/megalogo) to visualize this diversity metric via logoplot for long sequences with customizable annotations for secondary structure and the corresponding WT residue for each column. For these logoplots, the height of the AA is inversely proportional to the number of total AAs at the site and does not indicate the relative frequencies of these amino acids in the virus library.

The filtered mutVirus.p2 dataset with the *ndet* metric (**Figure 1E**) was passed through *megaLogo* and plotted with the WT CHIKV AF15561 reference sequence annotated above each column (**Figure 2A**). We then generated a heatmap for the corresponding *ndet* values for each residue in the trimeric surface glycoprotein (PDB: 3J2W) as well as the p62/E1 complex (PDB: 3N42) to visualize the mutational tolerance of E3 (given the absence of E3 from available CHIKV cryo-EM trimer structures) (**Figure 2B**). The logoplot and structural heatmaps for mutVirus.p2 (**Figures 2A-B**) revealed several individual sites (e.g., WT cysteine residues shown in **Figure 2A**) and conformationally relevant regions (e.g., the trimer core and center ridge within the E2 B domain shown in **Figure 2B**) where limited mutational tolerance was observed, suggesting these sites serve important purposes for protein-protein interactions and/or the virus’ life cycle. This is in contrast to more mutationally-tolerant spans, such as within E3 (e.g., positions 13-25), and within both the E2 A and B domains, evidenced by larger stacks of residues in **Figure 2A** and many residues with darker red shading in **Figure 2B**. Plotting the distribution of *ndet* per site for each domain revealed an increased number of sites in the E2 B domain with higher numbers of detected AAs when compared with other E2 domains (**Figure 2C**), suggesting the B domain has more plasticity in sites where mutations could be acquired before compromising viral fitness (45).

**Figure 2.**
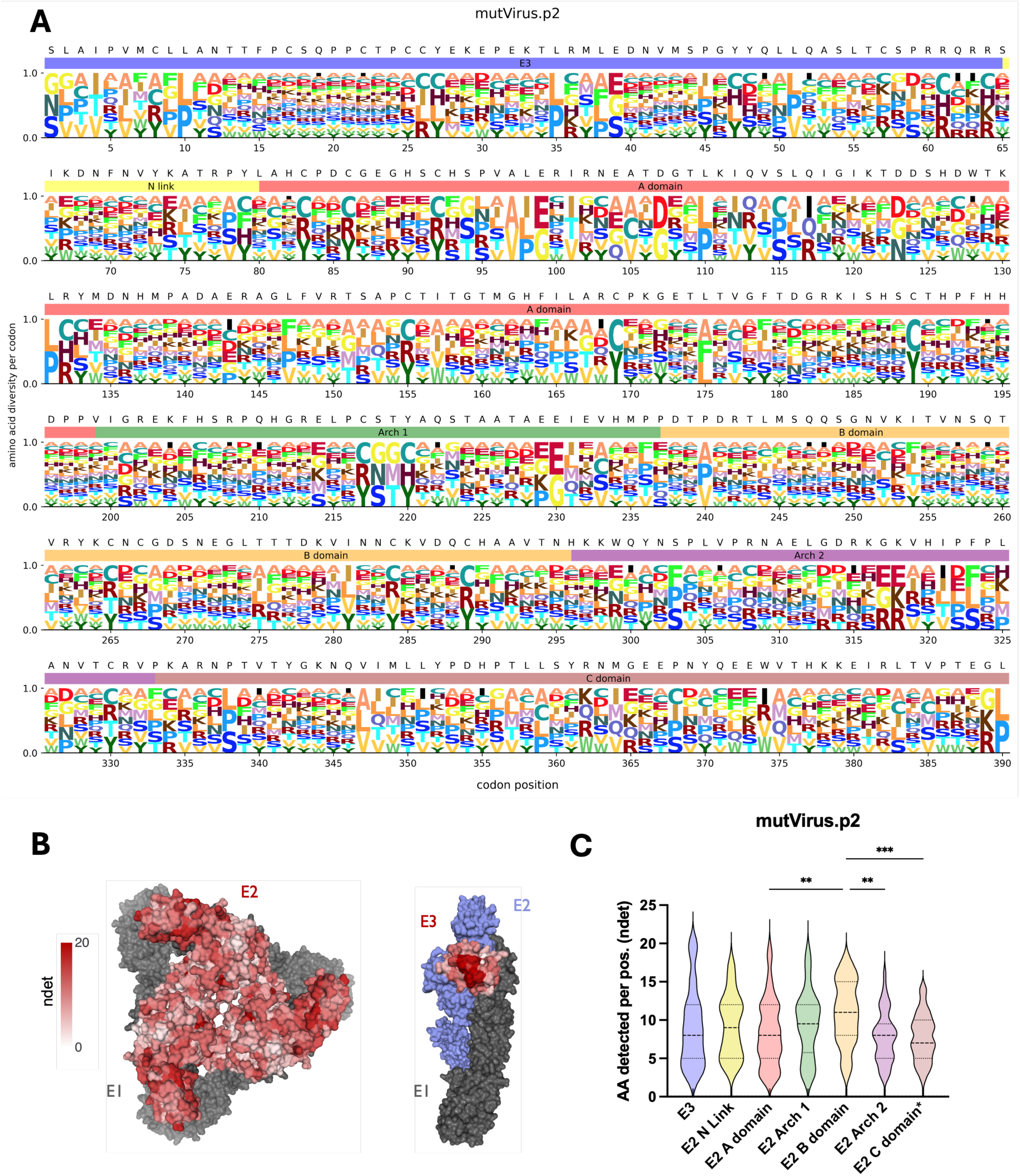
Deep mutational scanning of CHIKV p62 reveals mutational tolerance of the different p62 domains. (**A**) Logoplot showing the diversity of amino acids per codon position for the mutVirus.p2 virus library. WT residues are shown above each codon position and colored bars for each p62 domain below (E3: light blue, E2 N Link: light yellow, E2 A domain: light red, E2 Arch 1: light green, E2 B domain: light orange, E2 Arch 2: light purple, E2 C domain: light brown). Size of each letter is normalized to the number of amino acids detected for that codon position. (**B**) Using *dms-viz* (61), heatmaps were generated for the trimeric E2/E1 CHIKV envelope glycoproteins cryo-em structure (PDB: 3J2W) and the mature envelope glycoprotein complex (p62/E1; PDB: 3N42). For the trimeric structure, the heatmap represents E2 diversity from a top-down view. For the p62/E1 complex, the heatmap represents E3 diversity from a side view (with E2 colored in blue to highlight E3). E1 is shown in gray. (**C**) Violin plots showing the relative diversity at each site of each mutagenized domain are plotted. Colors are matched to colors shown in logoplot annotations in Panel 2A. *The E2 C domain region only includes the ectodomain portion of the C domain. One-way ANOVA with Tukey’s test for multiple comparisons. ** p< 0.01, *** p< 0.001.

### Mapping of Ab Escape for CHIKV nAbs Identifies New Sites of Escape

To validate the use of the CHIKV-p62-DMS library virus to identify mechanisms of viral escape from nAbs, we performed escape mutant selection assays with two well-characterized anti-CHIKV mAbs, CHK-152 and CHK-265, as well as an unmapped anti-CHIKV mAb, CHK-11 (21–26). Each mAb was pre-incubated at 37°C for 1 h with either wtVirus or mutVirus.p2 (MOI = 1 PFU/cell) with 2X EC_97_ mAb and virus-mAb mixtures were inoculated onto Vero cells, and the resulting virus output at 24 hpi was sequenced for differential selection. To account for the highest potential variability in the selection experiment, technical duplicates for all wells were performed and sequenced (mAb/virus and virus only wells) with each mAb/virus replicate compared to each virus only replicate and the final average score was calculated from all comparisons. These results showed modest positive site escape scores throughout p62 for all mAbs, however, individual mutants at several sites were found with high differential selection scores (>10 log_2_) (**Figure S2A-B**).

To validate our findings, we evaluated the extent to which positive escape sites were identified at known contacts sites or critical residues for the previously studied mAbs CHK-152 and CHK-265 (**Tables S1-2**) (21–26). Defined as having an average (log_2_) positive differential selection score ≥ 0.1 across all comparisons, 9/9 (100%) of prior published sites had at least one escape mutant for CHK-152 and 54/59 (92%) for CHK-265 (**Figure S3**), indicating consistency with prior studies. Total positive differential selection scores (log_2_) for each site were plotted as a heatmap on the CHIKV trimeric structure and the E3-focused p62/E1 heterodimer for CHK-152 (**Figure 3A**; **purple**), CHK-265 (**Figure 4A**; **orange**), and CHK-11 (**Figure 5A**; **green**).

**Figure 3.**
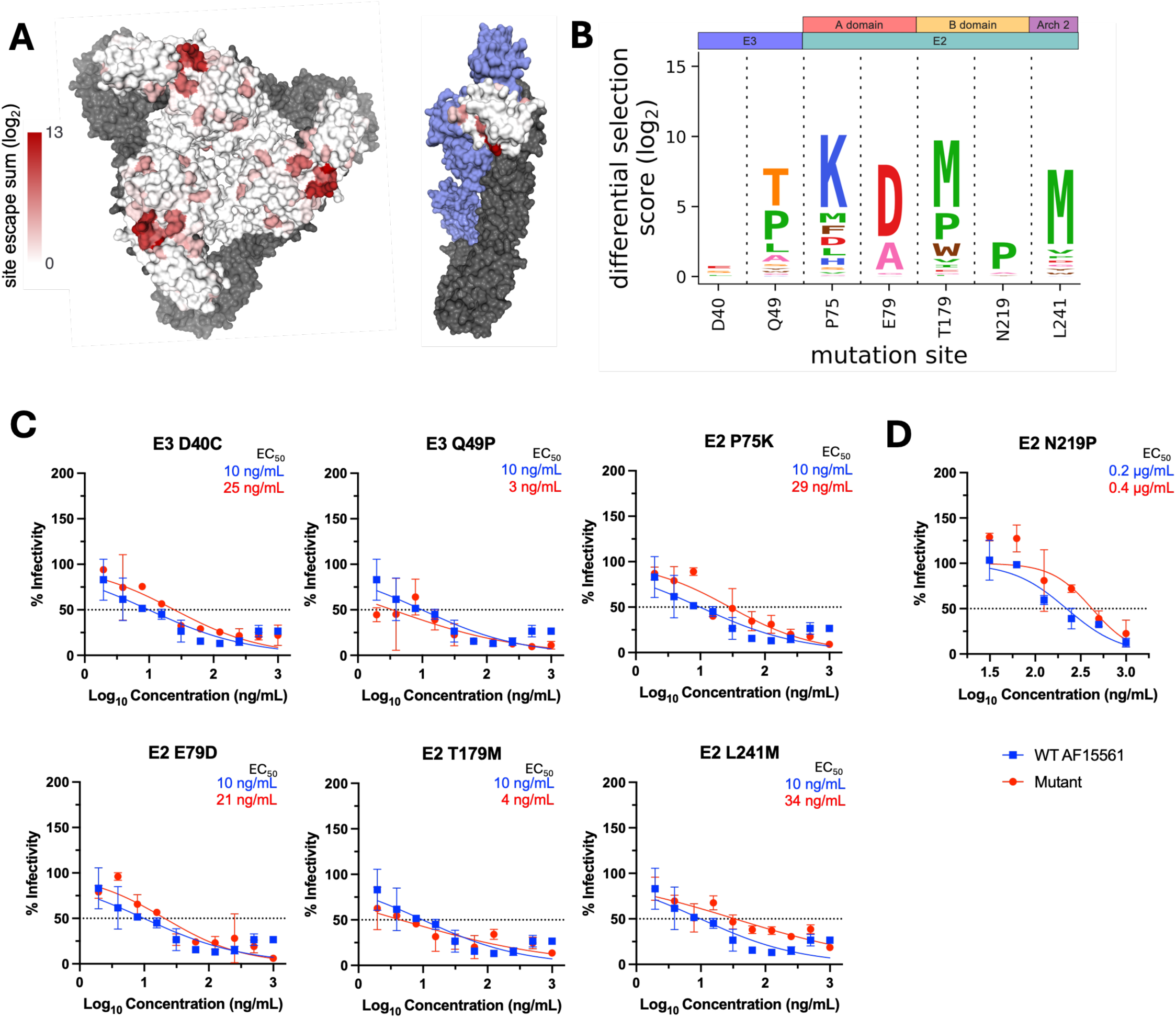
Escape mutant profile for CHK-152 monoclonal antibody reveals modest escape from selected panel of positive selection mutants. (**A**) Total site positive differential selection scores for CHK-152 were plotted via heatmap on the trimeric E2/E1 CHIKV envelope glycoproteins (PDB: 3J2W) and the mature envelope glycoprotein complex (p62/E1; PDB: 3N42). For the trimeric structure, the heatmap represents E2 positive site selection from a top-down view. For the p62/E1 complex, the heatmap represents E3 positive site selection from a side view (with E2 colored in blue to highlight E3). E1 is shown in gray for both structures. (**B**) Sites selected for validation of their sensitivity to neutralization by CHK-152. (**C**) FRNT curves for CHK-152 against WT CHIKV and the indicated mutant virus. Dotted line represents the FRNT_50_ threshold. (**D**) PRNT curve for CHK-152 against WT and E2 N219P CHIKV. Dotted line represents the PRNT_50_ threshold.

**Figure 4.**
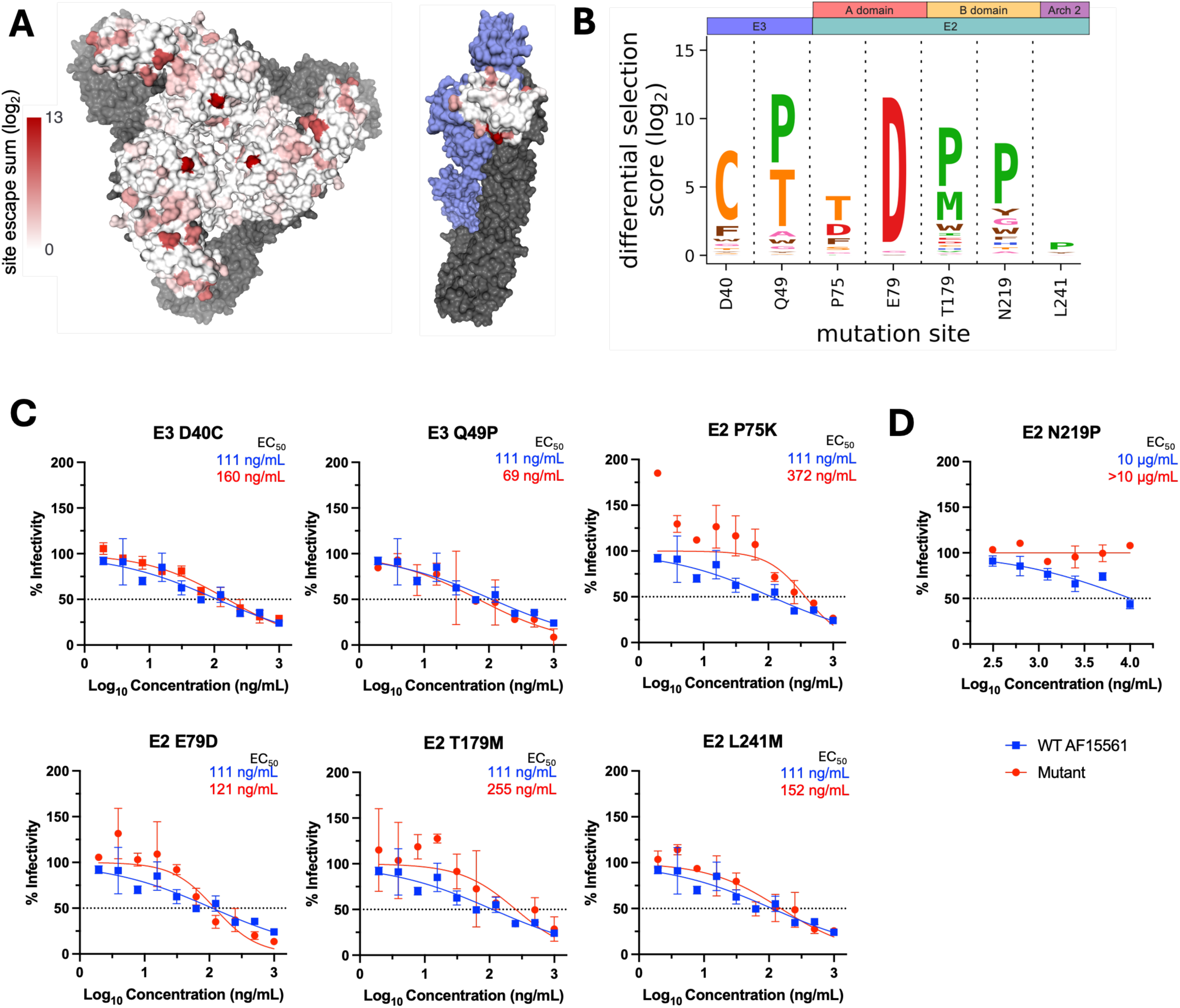
Escape mutant profile for CHK-265 monoclonal antibody reveals modest escape from selected panel of positive selection mutants. (**A**) Total site positive differential selection scores for CHK-265 were plotted via heatmap on the trimeric E2/E1 CHIKV envelope glycoproteins (PDB: 3J2W) and the mature envelope glycoprotein complex (p62/E1; PDB: 3N42). For the trimeric structure, the heatmap represents E2 positive site selection from a top-down view. For the p62/E1 complex, the heatmap represents E3 positive site selection from a side view (with E2 colored in blue to highlight E3). E1 is shown in gray for both structures. (**B**) Sites selected for validation of their sensitivity to neutralization by CHK-265. (**C**) FRNT curves for CHK-265 against WT CHIKV and the indicated mutant virus. Dotted line represents the FRNT_50_ threshold. (**D**) PRNT curve for CHK-265 against WT and E2 N219P CHIKV. Dotted line represents the PRNT_50_ threshold.

**Figure 5.**
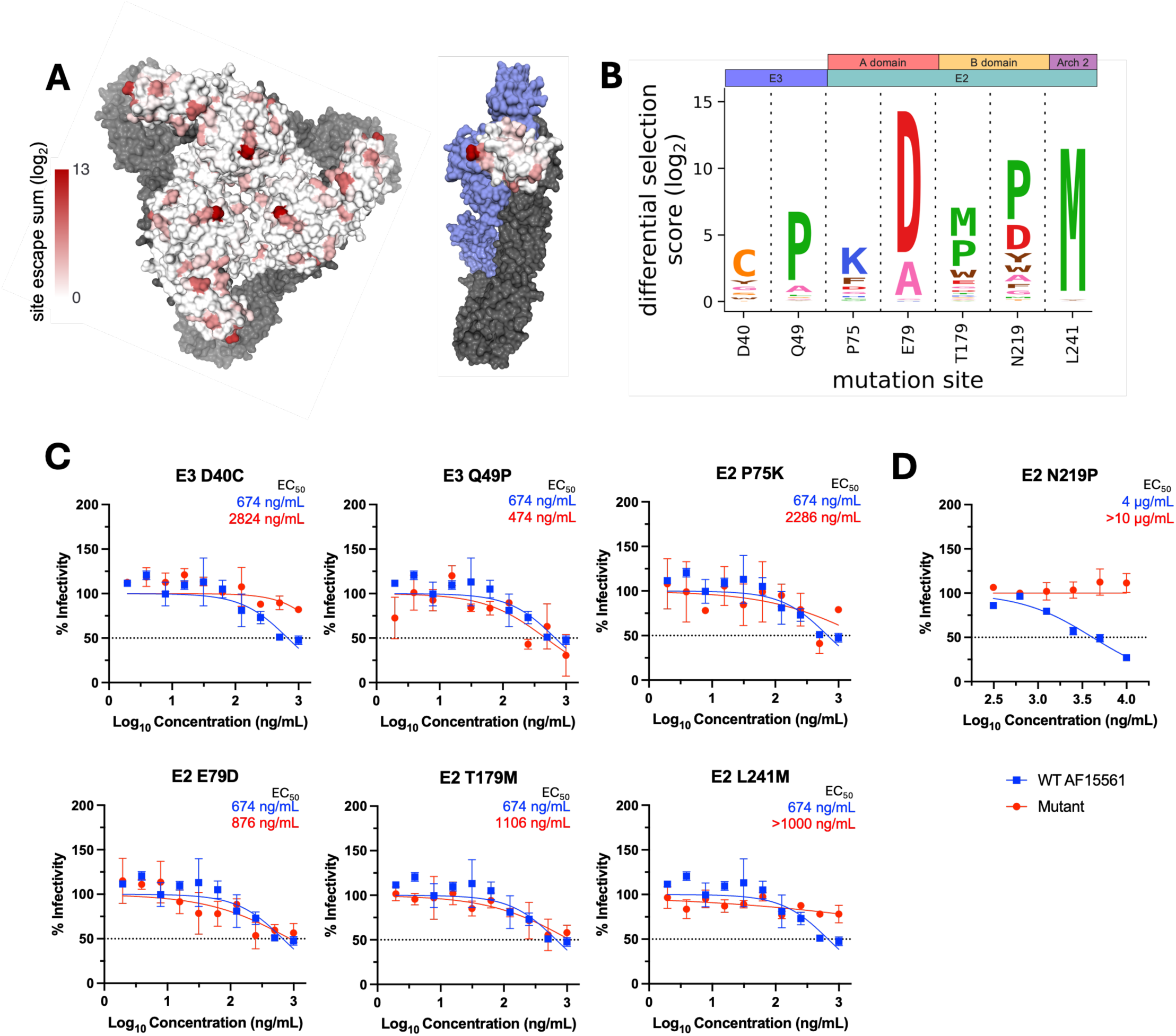
Escape mutant profile for CHK-11 monoclonal antibody reveals modest escape from selected panel of positive selection mutants. (**A**) Total site positive differential selection scores for CHK-11 were plotted via heatmap on the trimeric E2/E1 CHIKV envelope glycoproteins (PDB: 3J2W) and the mature envelope glycoprotein complex (p62/E1; PDB: 3N42). For the trimeric structure, the heatmap represents E2 positive site selection from a top-down view. For the p62/E1 complex, the heatmap represents E3 positive site selection from a side view (with E2 colored in blue to highlight E3). E1 is shown in gray for both structures. (**B**) Sites selected for validation of their sensitivity to neutralization by CHK-11. (**C**) FRNT curves for CHK-11 against WT CHIKV and the indicated mutant virus. Dotted line represents the FRNT_50_ threshold. (**D**) PRNT curve for CHK-11 against WT and E2 N219P CHIKV. Dotted line represents the PRNT_50_ threshold.

To determine if the CHIKV-p62-DMS library virus could be used to identify novel sites critical for Ab-mediated inhibition, we selected a panel of representative mutants at sites outside the known residues described in **Tables S1-2** to test for neutralization escape (**Figures 3-5**). In addition, we also tested the extent to which the CHIKV-p62-DMS library virus could be used to map escape from uncharacterized mAbs such as CHK-11. The criteria for selection of these sites included: (1) represent domains of interest with 1-2 sites, (2) score as a top escape mutant for at least two of the three mAbs, (3) not a previously identified site of escape, and (4) preferably surface exposed. This criterion produced the following mutants: E3 D40C, E3 Q49P, E2 P75K and E2 E79D (A domain), E2 T179M and E2 N219P (B domain), and E2 L241M (Arch 2) and the respective positive site escape logoplots (generated via *dmslogo*) are shown for each mAb (**Figures 3B, 4B**, **and 5B**).

We produced the panel of individual mutants in CHIKV AF15561 and evaluated neutralization capacity for each mAb (including an isotype negative control, anti-DENV nAb 2H2). Of note, one mutant virus, E2 N219P, was undetectable by our focus formation assay, which relies on CHK-11 as the detection Ab. For this reason, E2 N219P neutralization escape was evaluated for by a plaque reduction neutralization test (PRNT) (**Figures 3D, 4D**, **and 5D**) while all other mutants were evaluated by our standard FRNT assay (24). For most mutants tested in the FRNT assay, only modest escape was observed against each mAb with a 2-4-fold change in EC_50_ (ng/mL) (**Figures 3C**, **4C, and 5C**), except for E2 L241M which we were unable to calculate an EC_50_ for CHK-11 at the dilutions tested (**Figure 5C**). Conversely, the E2 N219P mutation mediated escape from mAbs CHK-265 and CHK-11 (**Figures 4D and 5D**) resulting in an inability to calculate an EC_50_ (µg/mL) for both mAbs. In contrast, E2 N219P mediated only modest 2-fold escape from CHK-152 (**Figure 3D**).

### CHK-11 Targets an Overlapping Epitope with CHK-265 and Mediates Broadly Neutralizing Activity Against Related Arthritogenic Alphaviruses

Due to the observed similarities between the escape profiles for CHK-11 and CHK-265 and neutralizing activity against the panel of escape mutants, we evaluated whether CHK-11 and CHK-265 target overlapping epitopes by competition ELISA (**Figure 6A**). Using an established whole virion based ELISA (24), biotinylated CHK-11 (BT CHK-11) was competed with unlabeled competitor Ab. Relative to BT CHK-11 binding with no competitor Ab, CHK-265 blocked binding of CHK-11 to a similar level as the self-competition control (CHK-11+BT CHK-11: 10.5%, CHK-265+BT CHK-11: 11%), suggesting significant epitope overlap or steric hinderance by CHK-265. CHK-152 only moderately reduced binding of CHK-11 (83.5%), suggesting more limited epitope overlap or steric hinderance (**Figure 6A**).

**Figure 6.**
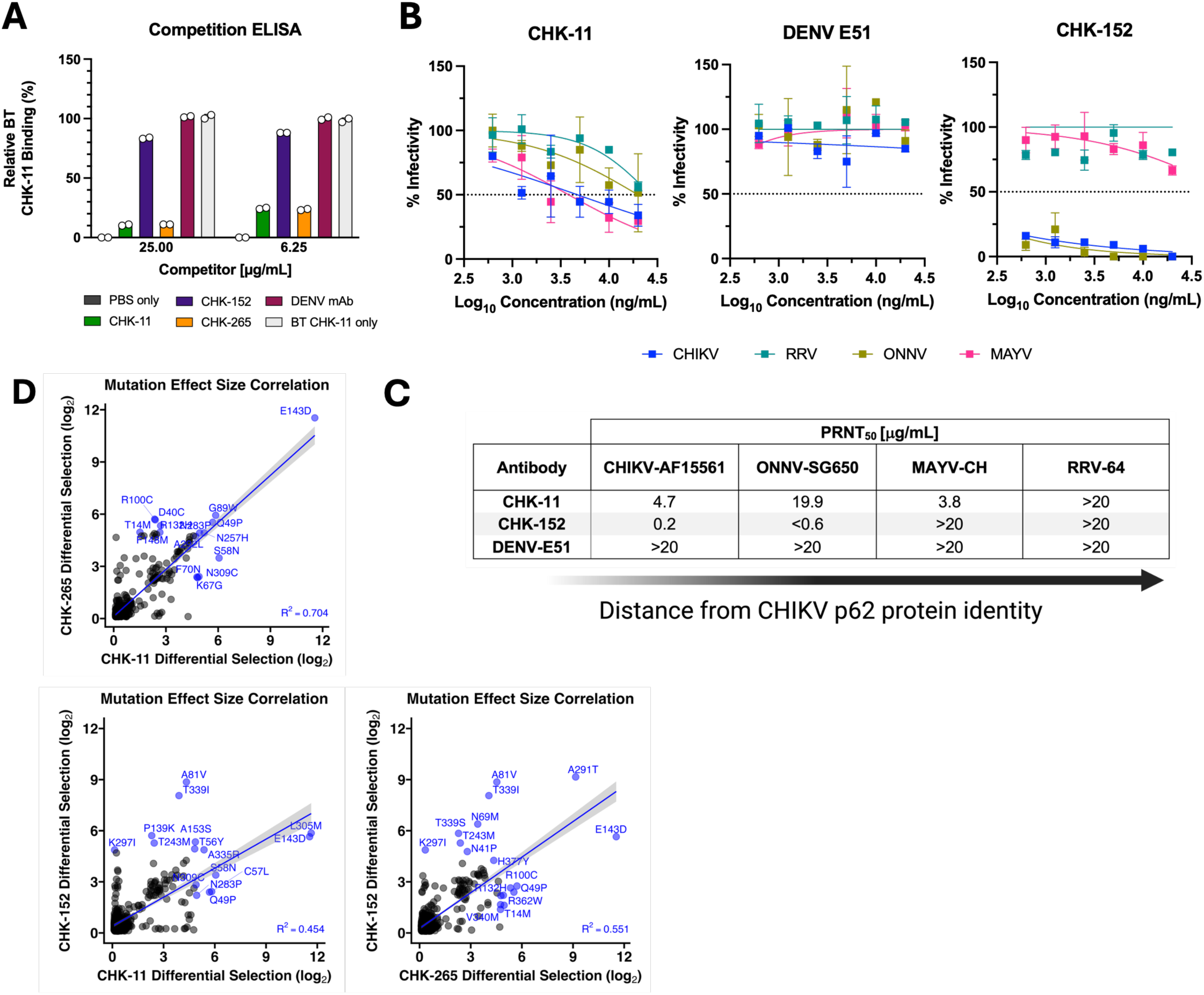
Validation of CHK-11 and CHK-265 similarity. (**A**) Competition ELISA against CHIKV virions (strain 181/25) with competitor antibodies added prior to the addition of biotinylated CHK-11. Percent binding (Absorbance @ 450 nm) relative to BT-CHK-11 alone (no competitor) was determined. (**B**) PRNT curves against CHIKV, Ross River virus (RRV), o’nyong’nyong virus (ONNV), and Mayaro virus (MAYV) on Vero cells. DENV E51 was included as an isotype, neutralizing antibody negative control. Dotted line represents the PRNT_50_ threshold. (**C**) Calculation of the PRNT_50_ in µg/mL for each virus-antibody condition, ordered by relatedness to CHIKV. (**D**) Correlation of shared positively selected mutants between CHK-11, CHK-152, and CHK-265 was evaluated to compare mutation effect sizes (log_2_ differential selection score) using linear regression in R v4.2.1. Top ten mutants for each mAb pair are colored blue.

Prior studies demonstrated that CHK-265 is a broadly neutralizing mAb against arthritogenic alphaviruses (23). Thus, we investigated if CHK-11 also shared this feature (21–26). Using the PRNT assay, we compared the neutralizing activity of CHK-11 and CHK-152, along with an isotype-matched control mAb (DENV-E51), against three additional arthritogenic alphaviruses, Ross River virus (RRV), o’nyong’nyong virus (ONNV), and Mayaro virus (MAYV) (**Figure 6B-C**). Similar to prior data published for CHK-265 (23, 26, 46), CHK-11 neutralized both CHIKV and MAYV, while neutralizing ONNV and RRV to a lesser degree. In contrast, CHK-152 neutralized both CHIKV and ONNV, while only minimal reductions in infectivity were noted for RRV and MAYV (**Figure 6B-C**).

Finally, to test if the observed similarities in their breadth and neutralization capacity of different alphaviruses extended back to our escape mutants identified for CHK-11 and CHK-265, we performed a correlation analysis of the differential selection scores (log_2_) for escape mutants shared by both mAbs (**Figure 6D**). Using a linear regression model analysis of all positive escape mutants identified in both datasets, we found a strong correlation between the average effect size of escape mutants for CHK-11 and CHK-265 (R^2^ = 0.704, p < 0.0001) with the top ten mutants for each mAb colored in blue (n = 15, five shared between antibodies). This contrasted with CHK-11 and CHK-152 (R^2^ = 0.454, p < 0.0001) and CHK-265 and CHK-152 (R^2^ = 0.551, p < 0.0001) which showed weaker correlations on differential selection scores for individual mutants, indicating CHK-11 and CHK-265 may share more contact sites or conformational dependencies than either antibody when evaluated against CHK-152.

## DISCUSSION

Outbreaks of CHIKV continue to grow in both size and frequency. Thus, it is important to define how the immune system (e.g., through Ab-mediated mechanisms) can be modulated to better protect against future outbreaks. Here, we describe the generation, characterization, and functional validation of a deep mutationally scanned CHIKV p62 virus library capable of rapidly characterizing the critical functional sites of several mAbs to address this need.

Using this library, we found that different regions of the CHIKV p62 ectodomain differentially tolerated the introduction of individual amino acids at each position, with the E2 B domain having a higher proportion of sites capable of tolerating more amino acid substitutions, specifically when compared to the E2 A, Arch 2, and C domains. This may be explained by the wide variety of protein interactions the E2 B domain can engage in, including both cell receptor and Ab binding, and this plasticity has been previously shown to permit rapid adaptation to restrictive cell types *in vitro* (47, 48). Conversely, mapping of the mutational tolerance at each site to the trimeric structure of CHIKV revealed several regions of the trimer with less lenience for substitutions, including portions of the surface-exposed region of the E2 A domain and trimer “core”, representing potential targets for therapeutics or vaccine immunogens. Additionally, the function of the E3 glycoprotein has not been fully elucidated. Prior studies have linked E3 to the virus maturation process (47), however, the mutational tolerance data this study provides should aid the field in identifying additional E3 functions, as well as the domains and residues critical for those functions.

Initially, we sought to verify the library’s capability to identify known functional sites (contact or functional escape) of well characterized mAbs. We chose two mAbs targeting different epitopes and confirmed that our mutagenized virus library identified previously published sites critical for each mAb, including specific functional escape mutants (i.e., E2 D59N for CHK-152 (21)). We also tested whether our comprehensive library could supplement the list of previously identified sites for these nAbs. A select panel of mutagenized viruses confirmed a modest escape mutant in E3, in addition to E2 A and B domain mutants. One mutant, E2 N219P, ablated the neutralization activity of CHK-265 which had been previously identified as an *in vivo* escape mutant during RRV infection (E2 T219P) following CHK-265 treatment (49) but not as a hit in traditional alanine mutagenesis scanning assays (23), further demonstrating the utility of more comprehensive, DMS-derived functional site mapping assays.

To utilize the virus library to functionally map candidate therapeutic mAbs for functional site-specificity in a high-throughput manner, we tested the capacity for the library to identify key residues for neutralization function of an unmapped mAb, CHK-11 (21). Using the same panel of mutants for CHK-11 as was used for CHK-152 and CHK-265, CHK-11 demonstrated a strong similarity to CHK-265 in sensitivity to certain mutants within the panel but was more sensitive to E3 D40C as well as the Arch 2 mutant, E2 L241M. A competition ELISA confirmed CHK-265 either overlaps or sterically hinders CHK-11 binding, and we found that CHK-11 shared the same broadly neutralizing activity profile as CHK-265 and other B domain bnAbs (23, 26, 49). This collection of data, along with correlation analyses with known mAbs CHK-152 and CHK-265, provide evidence the CHIKV-p62-DMS virus library and neutralization escape assay can be used to map critical Ab sites within the p62 ectodomain and can be further explored for its utility in mapping functional residues of polyclonal serum or additional mAbs in future studies.

This study has limitations. With the method used to introduce mutations throughout the p62 region, acquiring multiple mutations per clone is a possibility, as shown in our Sanger sequencing results. Additionally, the mutagenized region length exceeds a standard read length on most short-read sequencing platforms. Thus, phenotypes that were observed by deep sequencing could be confounded by mutations outside our read frames. To combat this, we verified if deep sequencing results accurately predicted escape mutants by individually introducing representative escape mutations and testing neutralization escape. This did confirm the phenotype expected for many of the mutants tested, however the degree of escape was more modest for some of the mutations than predicted by sequencing. However, especially for the bnAbs tested, CHK-265 and CHK-11, this may be due to the nature of their broadly neutralizing activity, as described in Kikawa et al (50) or the ability of bnAbs to inhibit egress in addition to blocking viral entry (23). Additionally, it is possible alphaviruses have virus-specific factors that contribute to the degree of differential selection observed when compared with prior DMS Ab site mapping studies that identified higher degrees of escape (31, 32). Future studies could incorporate a larger panel of Abs and escape mutants to determine if our findings extend to other anti-CHIKV mAbs or if Abs narrower than CHK-11/CHK-265 and even CHK-152 would have larger site differential selection scores within linear spans of residues, versus very large individual mutant scores throughout the mutagenized region.

Overall, this study demonstrates a method for characterizing alphavirus mutational tolerance and can be used to map critical sites for nAbs in a high-throughput system. Broadly, this research could provide insights into how the adaptive immune response can be modulated to better protect against infection and disease from CHIKV and other emerging alphaviruses.

## METHODS

### Plasmid Design

The CHIKV AF15561 sequence (GenBank accession no. EF452493) was cloned downstream of a human CMV promoter. An SV40 poly A sequence was inserted downstream of the CHIKV 3′-UTR and the hepatitis delta (HDV) ribozyme was inserted adjacent to the poly A tail of the viral genome. To enable identification of infected cells and as an attenuated biosafety feature (51–53), the viral genome was engineered to encode the fluorescent protein mKate in-frame in the viral structural polyprotein (54). ApaI and XhoI sites were also introduced, flanking the CHIKV p62 ectodomain to facilitate cloning of the mutagenized p62 fragments into the pCHIKV-CMV-mKate backbone. Two additional variations of this plasmid were produced, 1) digested with ClaI and religated to remove the CHIKV nonstructural proteins and prevent background caused by template carry-over during library mutagenesis, and 2) two stop codons were engineered into the E3 portion of the mutagenized p62 region to reduce potential overrepresentation of WT pCHIKV-CMV-mKate plasmids that were either single-cut or uncut during restriction enzyme digest.

### Library Mutagenesis and Cloning

The pCHIKV-CMV-mKate plasmid was digested with ClaI and re-ligated to isolate the p62-containing fragment and reduce the risk for WT virus carry-over. The resulting plasmid was the template to amplify the E3/E2 glycoprotein region for mutagenesis. PCRs were performed as previously described with slight modifications (37, 55). Primers for mutagenesis were designed using the NNK approach and generated using the *Codon Tiling Primers* Python script that prioritizes equal melting temperatures for oligos over equal-length oligos, then pooled all primers at equimolar concentration (31, 56). Forward and reverse mutagenesis PCRs, as well as the joining reaction PCRs, were performed for a single round to reduce codon mutation frequency. The joining reaction PCR product was gel-purified using the Zymoclean DNA Gel Recovery Kit (ZymoResearch). The mutagenized E3/E2 region and the pCHIKV-CMV-mKate plasmid (with two subsequent stop codon mutations in E2) were digested with ApaI and XhoI (NEB). Following dephosphorylation using alkaline phosphatase (NEB), ligation was performed using T4 DNA Ligase (NEB).

The ligation reaction product was electroporated into ElectroMAX^TM^ DH10B *E. coli* cells (ThermoFisher) to generate the plasmid library. Transformation was performed using 0.1 cm cuvettes (BioRad) at 2.0 kV, 200 W, 25 mF then incubated in SOC for 1 h at 37°C and plated on to 2X LB-Amp plates and incubated at 37°C for 14-18 h. Individual colonies were selected, grown in 2X LB-Amp broth, and miniprepped for analysis by Sangar sequencing. The remaining plated colonies were scraped and grown in 2X LB-Amp broth overnight and maxiprepped the following day. The total number of colonies was estimated by counting plates with either 1:40 or 1:400 dilutions of transformed bacteria.

### HEK293 Plasmid Library Transfection and Library Virus Infection

Mutant virus libraries were generated by transfection of HEK293 cells (ATCC: CRL-1573). Cells were plated for transfection with 30 μg mutDNA in 150 mm TC-treated dishes with 1.0 x 10^7^ cells and incubated overnight at 37°C prior to transfection. DNA transfection was performed using Lipofectamine 3000 (ThermoFisher) per manufacturer recommendations. Virus supernatants and cells were collected at 48 hpt.

Library virus infection was performed in 150 mm TC-treated dishes plated with 1.0 x 10^7^ Vero cells. Virus (MOI of 0.01 PFU/cell) was adsorbed to cells for 1 h at 37°C. After the adsorption period, D10 media was added, and cells were incubated at 37°C for 48 h.

### Virus Titering Assays

Focus formation assays (FFA) were performed as previously described (57). Foci were counted using the CTL BioSpot analyzer and software (Cellular Technology). FFA titers were calculated as the number of foci per milliliter of viral supernatant. For ONNV, MAYV, RRV, and CHIKV E2 N219P (and WT CHIKV control virus), titer assays were performed using a Plaque Formation Assay as described previously (57). Data was analyzed using Microsoft Excel and GraphPad Prism v10.1.1.

### Hybridoma Growth, Harvest and Purification of Monoclonal Antibodies (mAbs)

Hybridomas for CHK-11, CHK-152, and CHK-265 were kindly provided by Michael Diamond (Washington University) (21). In brief, hybridomas were expanded in complete IMDM media (Gibco; IMDM, 10% FBS, 1% penicillin-streptomycin) and transferred to a 2 L roller bottle at a density of 5 x 10^5^ cells/mL with 500 mL of IMDM and incubated in a 37°C roller incubator for 7 days. At 1 week, cells were supplemented with an additional 500 mL of complete IMDM without FBS. Cells were pelleted at 4°C and harvested when most cells were dead (>90%). Supernatant was filter sterilized and concentrated using Centricon Plus-70 100 kDa filters (Millipore Sigma). Concentrated supernatants were purified with the Nab Protein G Spin Kit (Life Technologies) and concentrated again using Amicon Ultra-15 10kda filters (Millipore Sigma). Final concentrates were quantified via a virion-based ELISA described previously (24).

### Plaque and Focus Reduction Neutralization Tests (PRNT/FRNT)

FRNTs were performed as previously described (21, 24). Prior to the addition of virus to Vero cells, virus was preincubated with a dilution series of mAb for 1 h at 37°C in a 96-well V-bottom plate along with virus-only controls. Following the addition of overlay at 2 hpi, cells were fixed with 2% PFA at 14-16 hpi. Data analysis was performed using Microsoft Excel and GraphPad Prism v10.1.1. The PRNT and plaque formation assay protocol is detailed in Hawman et al., with modification for a dilution series of monoclonal antibodies (as performed in the FRNT assay) instead of serum (with heat inactivation) (12, 24).

### Monoclonal Antibody Challenge

MAbs were incubated with either mutVirus.p2 or wtVirus.p2 at a virus MOI of 1 in 100 μL of diluent for 1 h at 37°C, and virus-mAb mixtures were added to Vero cells. Plates were incubated at 37°C for 1 h, cells were washed five times with 1X PBS, and 1 mL of fresh medium was added. At 16-18 hpi, supernatants were harvested and stored at -80°C. Viral titers and viral RNA were harvested and sequenced as described above.

### Mutant Virus Production

Site-directed mutagenesis (SDM) of plasmid DNA was performed using the QuikChange II XL SDM Kit (Agilent Technologies) according to the manufacturer’s instructions, transformed into XL-10 Gold cells, miniprepped, and then sequence confirmed. Select clones were midiprepped (ZymoResearch) and prepared for in-vitro transcription.

In vitro transcription was performed as previously described (57). In brief, plasmids were linearized with NotI (NEB). *In vitro* transcription was performed using mMessage mMachine Kit (Invitrogen) per manufacturer instructions. BHK-21 cells (ATCC: CCL-10) were electroporated as previously described (58) and harvested 24 h later. Viral titers were determined by FFA.

### Verification of Critical Sites for CHK-152 and CHK-265

Literature review results for critical sites previously identified for neutralizing antibody CHK-152. Given all sites have been identified in E2 are detailed in **Tables S1-2**, E2 position numbering was included along with the respective p62 numbering for referencing any results described within this manuscript. If the strain in the study differed from our use of CHIKV AF15561, that was noted. The origin strain residue for that site was also included and if the amino acid differed in the AF15561 strain, the AF15561 residue was denoted as (X). In the case the study identified a mutation that escaped the tested function, this was recorded. All identified escape mutants in our study were listed in the “Library Escape Mutants” column in order of largest (log_2_) positive differential selection score to least (log_2_) positive differential selection score. The method the study used to identify the critical residue was also listed and referenced. Mutants with average (log_2_) positive differential selection scores ≥ 0.1 across all comparisons were included and ranked based on their average score and bolded if identified as an escape mutant in the referenced studies.

### RNA Isolation, RT-PCR, and Amplicon PCR for Library Preparation

RNA isolation of viral samples and RT-PCR for generation of cDNA was performed as described previously (59). To improve sequencing coverage of the mutagenized p62 region, an amplicon PCR was performed using KOD Hot Start (Millipore Sigma) as previously published (55) utilizing the amplifying primers described below (see ***Primers***). PCR amplicons were PCR-purified using QIAquick PCR Purification Kit (Qiagen) and submitted for mechanical shearing (Covaris) and library preparation to the University of Colorado Anschutz Medical Campus’ Genomics Shared Resource Facility (RRID: SCR_021984). Library preparation was performed using the Ovation Ultralow System V2 kit (Tecan).

### Sequence Analysis

All samples were sequenced with 2x150bp reads at a minimum depth of 25M/50M PE reads on the Illumina NovaSeq 6000. Raw fastq reads from the NovaSeq 6000 were evaluated for sequencing quality using *FastQC* software v0.11.9. Reads were then trimmed and cleaned using *cutadapt* software v3.4. The quality of trimmed reads was re-evaluated for quality using *FastQC*. These reads were then subsampled with *seqtk* to 25M PE reads and aligned to the WT pCHIKV-CMV-mKate plasmid reference genome (see **Data and Code Availability** section) and codons called using *VirVarSeq* software (60). The *VirVarSeq* output was filtered for mutations with a minimum average Phred quality score of 24 or higher and analyzed using a custom code in Python v3.9.2 and R v4.2.1. (see **Data and Code Availability** section). Amino acid preferences were calculated using *dms_tools2* (56). Structures were pulled from the Protein Data Bank (PDB: 3N42, 3J2W) and heatmap projections were generated using *dms-viz* (*61*). Bar plots, violin plots, and neutralization curves were plotted and analyzed using GraphPad Prism v10.1.1. Miniprep clone sequences were aligned to the pCHIKV-CMV-mKate reference genome using Geneious Prime v2022.1.1.

To calculate the *ndet* metric, we first filtered the *VirVarSeq* output codon matrices with the following conditions: (1) must have greater than 100 counts for that codon-position pair, and (2) must have an average forward and reverse read minimum Phred quality score ≥ 24. Then, the remaining unique residues were summed for each position (*ndet*). For plotting the filtered residues into logoplots, each position stack was set to total a value of 1.0 and the height of the AA (*F_site_*) is inversely proportional to the number of total AAs at the site (1/*nted*). Thus, the size of each residue does not indicate the relative frequencies of these amino acids in the virus library but rather the mutational tolerance at that codon position.

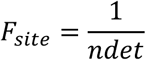

### Data and Code Availability

The deep sequencing data are available on the Sequence Read Archive under accession numbers SRA and will be available upon publication. All code used to analyze data included here and any remaining software outputs not published in this manuscript, along with reference sequence files, are available in the supplementary materials.

### Primers

The PCR primers used for both amplification and joining reactions were as follows: FwdPrimer: 5’-AGACGTTGAGTCCAACCCTGGGCCCA-3 and RevPrimer: 5’-CTCGTTGTTGCCCCACGTGACCTCGAG-3’. The list of NNK degenerative primers generated for the mutagenesis reaction is available upon publication at: https://github.com/meganstumpf/chikvdms-mAb-paper/. For Sangar sequencing of mutagenized miniprep clones, the following primers were used to cover the p62 ectodomain region with double coverage: CHIKV_Lib_FwdSeq1: 5’-CAAGGAGGCCGACAAAGAGAC-3’, CHIKV_Lib_FwdSeq3: 5’-CCAGGTTTCCTTGCAAATCGG-3’, CHIKV_Lib_RevSeq1: 5’-GCTAGGTACGGTCTTGTGGC-3’,

CHIKV_Lib_RevSeq2: 5’-CCACCGTCAGAGTTTCTCC-3’, CHIKV_Lib_RevSeq5: 5’-CAGGAGTACGAACGAGGCC-3’.

## ACKNOWLEDGEMENTS

We would like to thank Dr. Michael S. Diamond for kindly providing hybridomas for the anti-CHIKV mAbs utilized in our study. The following reagent was obtained through BEI Resources, NIAID, NIH: Monoclonal Anti-Dengue Virus Type 1 Envelope Protein, Clone E51 (produced *in vitro*), NR-4757. This work was supported by NIH grants R01 AI141436 and R01 AI148144 to T.E.M. This study was supported in part by the National Institutes of Health P30CA06934 funded Genomics Shared Resource [RRID:SCR_021984].

## AUTHOR CONTRIBUTIONS

**MMS:** conceptualization, data curation, formal analysis, investigation, methodology, software, visualization, writing – original draft, writing – review & editing

**TB:** data curation, formal analysis, resources, software, supervision, writing – review & editing

**MM:** conceptualization, data curation, methodology, writing – review & editing

**BJD:** methodology, supervision, writing – review & editing

**TEM:** conceptualization, data curation, funding acquisition, investigation, methodology, project administration, resources, supervision, visualization, writing – original draft, writing – review & editing

