## Supplemental Materials for "Deep mutationally scanned (DMS) CHIKV E3/E2 virus library maps viral amino acid preferences and predicts viral escape mutants of neutralizing CHIKV antibodies"

### SUPPLEMENTARY MATERIALS

#### *Data and Code Availability*

Raw FASTQ files from sequencing will be available to the public on GEO with accession numbers provided upon manuscript publication. Complete documentation of all analyses utilized in this manuscript will be available to the public on GitHub at https://github.com/meganstumpf/chikvdms-mAb-paper upon manuscript publication. Contact the corresponding author for information requests prior to manuscript publication.

*Supplementary Figures*

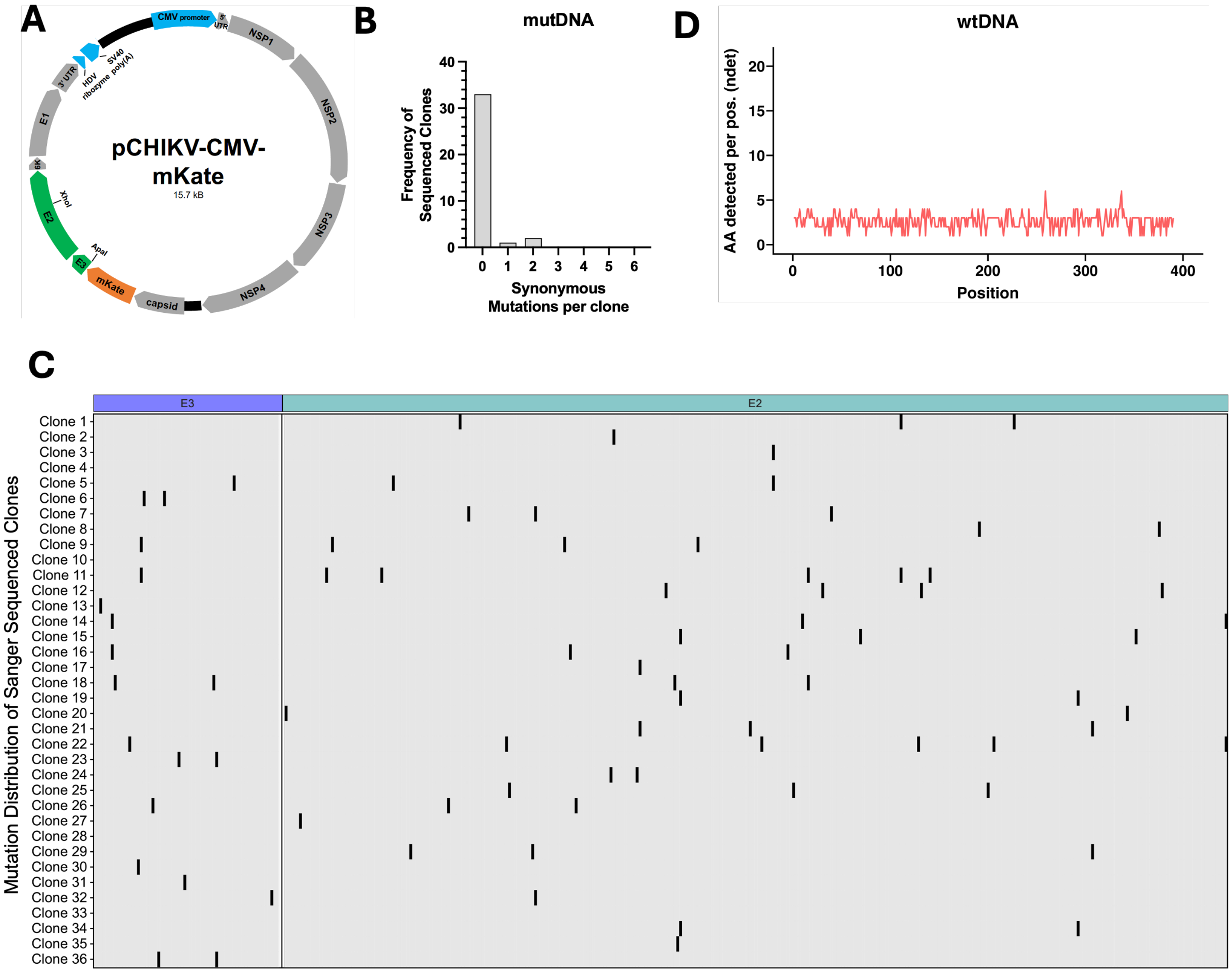


**Figure S1. Distribution of clonal mutations across representative mutDNA minipreps and diversity in wtDNA sequencing control.** (**A**) Plasmid design for pCHIKV-CMV-mKate. (**B**) Sanger sequencing results for individual plasmid clones and number of synonymous mutations per full-length CHIKV p62 clone (Sanger primers available in **Materials and Methods**. (**C**) Forty plasmid clones were Sanger sequenced and aligned to the pCHIKV-CMV-mKate WT control for number of amino acid mutations in the entire mutagenized p62 region. E3 is annotated by the periwinkle box on the left, E2 in teal on the right. Each black line represents a nonsynonymous mutation from the WT sequence. Four of the 40 clones (10%) were excluded from the alignment because they either represented vector-only clones or were non-CHIKV plasmid contaminants. Alignment performed using Geneious and graphed with GraphPad Prism. (**D**) The total number of amino acids detected per codon position ('ndet') for the wtDNA sequencing control. For all differential selection analyses, the wtDNA control is subtracted from all experimental conditions.

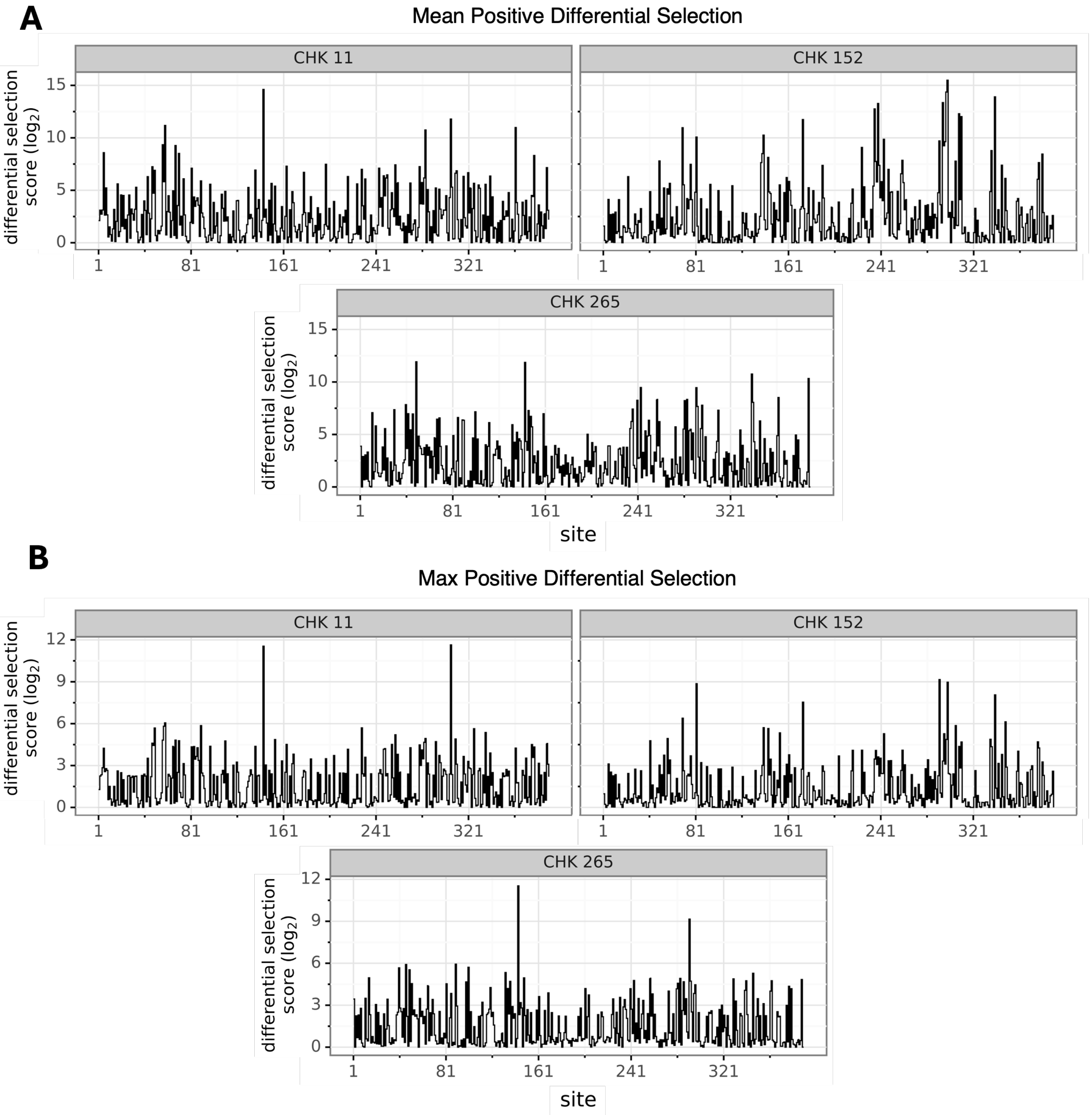


**Figure S2. Differential selection line plots for monoclonal antibodies CHK-11, CHK-152, and CHK-265 against the CHIKV-p62-DMS virus library.** Using *dms_tools2*, the average differential selection across replicate wells was calculated for each neutralizing antibody against both replicates of virus-only control wells independently and error corrected with the corresponding wtDNA sequencing control. (**A**) For all comparisons, the average positive differential site selection scores are plotted (measured as $\log_2$ scores) for each antibody. (**B**) The highest-scoring positively-selected mutant (measured as $\log_2$ scores) is plotted for each site after averaging across all comparisons for each antibody.

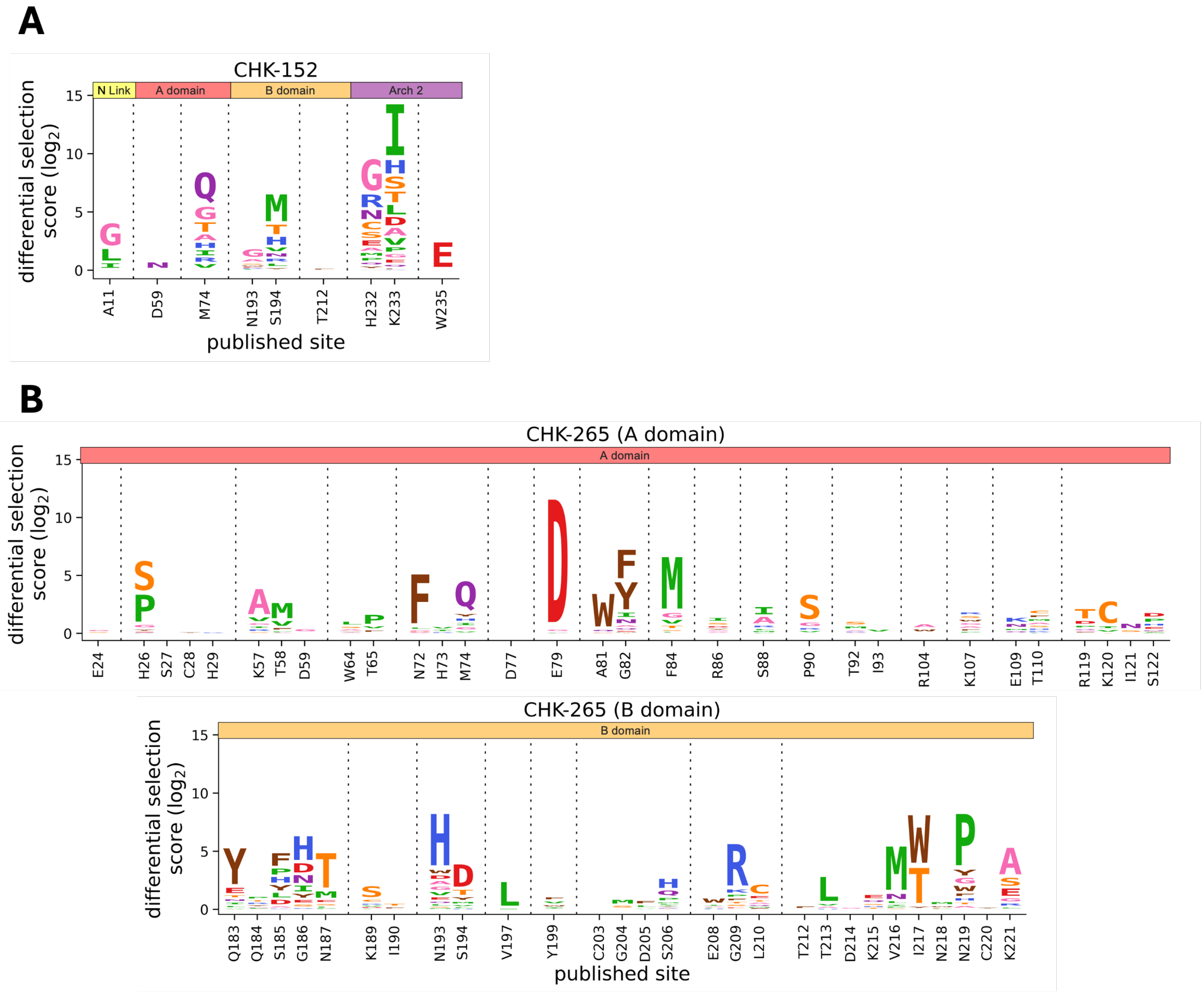


**Figure S3. Escape mutations identified at sites previously reported as contacts or critical residues for CHK-152 and CHK-265 monoclonal antibodies.** (**A and B**) Previously identified contact sites or other critical residues for antibodies CHK-152 (A) and CHK-265 (B) were recorded (**see Tables S1 and S2**). These sites were extracted from the differential selection dataset following escape mutant selection and positively selected mutants plotted via logoplot. Antibody escape is reported as a $\log_2$ selection score with letter heights corresponding to degree of selection for that mutant averaged across all comparisons with error correction from the wtDNA sequencing control. For (B), due to the large number of important sites identified for CHK-265, the logoplot was split by whether the site resides in the E2 A domain or E2 B domain, respectively.

| | Domain | E2 no. | p62 no. | Origin Strain | Origin Residue | Escape Residue | Library Escape Mutants† | Residue Mapping Method |
|---|---|---|---|---|---|---|---|---|
| **CHK-152** | **A** | 11 | 75 | CHIKV-LR | A | NA | G, L, I | Cryo-EM road mapping[1] |
| | | 59 | 123 | CHIKV-LR | D | N | **N** | Serial passaging[2] |
| | | | | | | NA | | Cryo-EM road mapping[1] |
| | | 74 | 138 | CHIKV-LR | M | NA | Q, G, T, A, H, I, R, V | Cryo-EM road mapping[1] |
| | **B** | 193 | 257 | CHIKV-LR | N | NA | G, A, S, W, E | Cryo-EM road mapping[1] |
| | | 194 | 258 | CHIKV-LR | G | NA | M, T, H, V, N, R, L, Y | Cryo-EM road mapping[1] |
| | | 212 | 276 | CHIKV-LR | T | NA | F | Cryo-EM road mapping[1] |
| | | 232 | 296 | CHIKV-LR | H | NA | G, R, N, C, S, E, A, M, P, Q, Y | Cryo-EM road mapping[1] |
| | **β-ribbon connector (Arch 2)** | 233 | 297 | rVSV-CHIKV | K | T | I, H, S, **T**, L, D, A, V, P, G, **E**, Q, F | In vivo, neutralization[2,3] |
| | | | | CHIKV-LR | | E | | In vivo selection[2] |
| | | 235 | 299 | CHIKV-LR | W | NA | E | Cryo-EM road mapping[1] |

**Table S1. Escape mutations identified at sites previously reported as contacts or critical residues for CHK-152 monoclonal antibody.**

| | Domain | E2 no. | p62 no. | Origin Strain | Origin Residue | Escape Residue | Library Escape Mutants† | Residue Mapping Method |
|---|---|---|---|---|---|---|---|---|
| CHK-265 | A | 24 | 88 | CHIKV-LR | E | NA | G, S | Cryo-EM road mapping[4] |
| | | 26 | 90 | CHIKV-LR | H | NA | S, P, G, Y | Cryo-EM road mapping[4] |
| | | 27 | 91 | CHIKV-LR | S | NA | none | Cryo-EM road mapping[4] |
| | | 28 | 92 | CHIKV-LR | C | NA | none | Cryo-EM road mapping[4] |
| | | 29 | 93 | CHIKV-LR | H | NA | none | Cryo-EM road mapping[4] |
| | | 57 | 121 | CHIKV-LR | K | NA | A, V, G, I, R | Cryo-EM road mapping[4] |
| | | 58 | 122 | CHIKV-LR | T | NA | M, V, F | Cryo-EM road mapping[4] |
| | | 59 | 123 | CHIKV-LR | D | NA | G | Cryo-EM road mapping[4] |
| | | 64 | 128 | CHIKV-LR | W | NA | L, S, A, Y | Cryo-EM road mapping[4] |
| | | 65 | 129 | CHIKV-LR | T | NA | P, V, F | Cryo-EM road mapping[4] |
| | | 72 | 136 | CHIKV-LR | N | NA | F, L, G | Cryo-EM road mapping[4] |
| | | 73 | 137 | CHIKV-LR | H | NA | V, I | Cryo-EM road mapping[4] |
| | | 74 | 138 | CHIKV-LR | M | NA | Q, Y, H, I, G, V | Cryo-EM road mapping[4] |
| | | 77 | 141 | CHIKV-LR | D | NA | none | Cryo-EM road mapping[4] |
| | | 79 | 143 | CHIKV-LR | E | NA | D, G | Cryo-EM road mapping[4] |
| | | 81 | 145 | CHIKV-LR | A | NA | W, Q | Cryo-EM road mapping[4] |
| | | 82 | 146 | CHIKV-LR | G | NA | F, Y, I, N, A, Q, E | Cryo-EM road mapping[4] |
| | | 84 | 148 | CHIKV-LR | F | NA | M, G, V, T, S, L | Cryo-EM road mapping[4] |
| | | 86 | 150 | CHIKV-LR | R | NA | I, S, D, Y, G, A | Cryo-EM road mapping[4] |
| | | 88 | 152 | CHIKV-LR | S | NA | I, A, R, L, M | Cryo-EM road mapping[4] |
| | | 90 | 154 | CHIKV-LR | P | NA | S, G, R, V | Cryo-EM road mapping[4] |
| | | 92 | 156 | CHIKV-LR | T | NA | S, M, A | Cryo-EM road mapping[4] |
| | | 93 | 157 | CHIKV-LR | I | NA | V | Cryo-EM road mapping[4] |
| | | 104 | 168 | CHIKV-LR | R | NA | A, W | Cryo-EM road mapping[4] |
| | | 107 | 171 | CHIKV-LR | K | NA | R, S, W, G, A, F, P | Cryo-EM road mapping[4] |
| | | 109 | 173 | CHIKV-LR | E | NA | K, N, M, Q | Cryo-EM road mapping[4] |
| | | 110 | 174 | CHIKV-LR | T | NA | C, F, M, A, S, H, K | Cryo-EM road mapping[4] |
| | | 119 | 183 | CHIKV-LR | R | NA | T, D, P, Q, G, F | Cryo-EM road mapping[4] |
| | | 120 | 184 | CHIKV-LR | K | NA | C, I, V | Cryo-EM road mapping[4] |
| | | 121 | 185 | CHIKV-LR | I | NA | N, S | Cryo-EM road mapping[4] |
| | | 122 | 186 | CHIKV-LR | S | NA | D, P, H, E | Cryo-EM road mapping[4] |
| | B | 183 | 247 | CHIKV-37997 | Q | NA | Y, E, T, N, I, L | PISA solvent exclusion analysis[5] |
| | | 184 | 248 | CHIKV-LR | Q | A | P, T, R, Y | Alanine-scanning mutagenesis[4] |
| | | | | | | NA | | Cryo-EM road mapping[4] |
| | | | | CHIKV-37997 | | NA | | PISA solvent exclusion analysis[5] |
| | | 185 | 249 | CHIKV-LR | S | A | F, P, H, Y, L, D, **A** | Alanine-scanning mutagenesis[4] |
| | | | | | | NA | | Cryo-EM road mapping[4] |
| | | | | CHIKV-37997 | | NA | | PISA solvent exclusion analysis + alanine scanning mutagenesis/neutralization escape[5] |
| | | 186 | 250 | CHIKV-37997 | G | NA | H, D, N, I, Y, E, S | PISA solvent exclusion analysis + alanine scanning mutagenesis/neutralization escape[5] |
| | | 187 | 251 | CHIKV-LR | N | NA | T, M, E, S, L | Cryo-EM road mapping[4] |
| | | | | CHIKV-37997 | | NA | | PISA solvent exclusion analysis[5] |
| | | 189 | 253 | CHIKV-LR | K | NA | S, C, R, V, D | Cryo-EM road mapping[4] |
| | | | | CHIKV-37997 | | NA | | PISA solvent exclusion analysis[5] |
| | | 190 | 254 | CHIKV-LR | I | A | T, F | Alanine-scanning mutagenesis[4] |
| | | 193 | 257 | CHIKV-LR | D (N) | NA | H, W, D, A, G, V, E, Q, L, S | Cryo-EM road mapping[4] |
| | | 194 | 258 | CHIKV-LR | E (S) | NA | D, T, Y, M, V, L | Cryo-EM road mapping[4] |
| | | 197 | 261 | CHIKV-LR | V | A | L, I | Alanine-scanning mutagenesis[4] |
| | | 199 | 263 | CHIKV-LR | Y | A | F, V, W, I | Alanine-scanning mutagenesis[4] |
| | | 203 | 267 | CHIKV-37997 | C | NA | none | PISA solvent exclusion analysis[5] |
| | | 204 | 268 | CHIKV-37997 | G | NA | M, S | PISA solvent exclusion analysis[5] |
| | | 205 | 269 | CHIKV-LR | G (D) | NA | F, L | Cryo-EM road mapping[4] |
| | | 206 | 270 | CHIKV-LR | S | NA | H, Q, P, M, L, I, Y | Cryo-EM road mapping[4] |
| | | 208 | 272 | CHIKV-LR | E | NA | W, I, A | Cryo-EM road mapping[4] |
| | | 209 | 273 | CHIKV-LR | G | A | R, K, P, F, T, S, E | Alanine-scanning mutagenesis[4] |
| | | 210 | 274 | CHIKV-LR | L | A | C, E, T, **A**, Q, M | Alanine-scanning mutagenesis[4] |
| | | | | | | NA | | Cryo-EM road mapping[4] |
| | | 212 | 276 | CHIKV-LR | T | A | F | Alanine-scanning mutagenesis[4] |
| | | | | | | NA | | Cryo-EM road mapping[4] |
| | | 213 | 277 | CHIKV-LR | T | NA | L, V, A | Cryo-EM road mapping[4] |
| | | 214 | 278 | CHIKV-LR | D | NA | G | Cryo-EM road mapping[4] |
| | | 215 | 279 | CHIKV-LR | K | NA | E, Q, S, H, T | Cryo-EM road mapping[4] |
| | | 216 | 280 | CHIKV-LR | V | NA | M, N, L, I, P | Cryo-EM road mapping[4] |
| | | | | CHIKV-37997 | | NA | | PISA solvent exclusion analysis[5] |
| | | 217 | 281 | CHIKV-LR | I | A | W, T, Y | Alanine-scanning mutagenesis[4] |
| | | 218 | 282 | CHIKV-LR | N | NA | M, W | Cryo-EM road mapping[4] |
| | | | | CHIKV-37997 | N | NA | | PISA solvent exclusion analysis[5] |
| | | 219 | 283 | CHIKV-LR | N | NA | P, Y, G, W, F, H, T, A | Cryo-EM road mapping[4] |
| | | | | CHIKV-37997 | | NA | | PISA solvent exclusion analysis[5] |
| | | 220 | 284 | CHIKV-37997 | C | NA | Y | PISA solvent exclusion analysis[5] |
| | | 221 | 285 | CHIKV-37997 | K | NA | A, S, E, G, R, L | PISA solvent exclusion analysis[5] |

**Table S2. Escape mutations identified at sites previously reported as contacts or critical residues for CHK-265 monoclonal antibody.**
